# Action boosts episodic memory encoding in humans via engagement of a noradrenergic system

**DOI:** 10.1101/203034

**Authors:** Mar Yebra, Ana Galarza-Vallejo, Vanesa Soto-Leon, Javier J Gonzalez-Rosa, Archy O de Berker, Sven Bestmann, Antonio Oliviero, Marijn CW Kroes, Bryan A Strange

## Abstract

We are constantly interacting with our environment whilst we encode memories. However, how actions influence memory formation remains poorly understood. Goal-directed movement engages the locus coeruleus (LC), the main source of noradrenaline in the brain. Noradrenaline is also known to enhance episodic encoding, suggesting that action could improve memory via LC engagement. Here we demonstrate, across seven experiments, that action (Go-response) enhances episodic encoding for stimuli unrelated to the action itself, compared to action inhibition (NoGo). Supporting a noradrenergic mechanism underlying this enhancement, functional magnetic resonance imaging, and pupil diameter as a proxy measure for LC-noradrenaline transmission, indicate increased encoding-related LC activity during action. A final experiment confirmed a novel prediction derived from these data that emotionally aversive stimuli, which recruit the noradrenergic system, modulate the mnemonic advantage conferred by Go-responses relative to neutral stimuli. We therefore provide converging evidence that action boosts episodic memory encoding via a noradrenergic mechanism.

---

Much of the episodic memories we form in daily life are encoded whilst we are physically active. However, the extent to which actions influence episodic memory encoding is currently unknown. Research on educational techniques shows that “active learning”, an instructional method that stimulates student activity in class, such as making button-press responses, is more effective than passively receiving information from an instructor^1,2^. However, within this framework, students typically make motor responses to questions posed by the instructor, thus the effect of the action *per se* (button-press alone) on learning cannot be identified. A separate line of study has shown that memory for action phrases (*e.g.*, “pick up the book”) is improved when participants perform the actions during encoding compared with merely listening to or reading the phrases^3,4^. Yet, since the memory tested in this task pertains to the movement, the memory of the movement cannot be dissociated from the effect of engaging the motor system on memory. In other words, it is currently unknown whether actions influence memory for stimuli that are incidental to the movement being carried out. An example of the latter effect would be whether the likelihood of remembering the title of a book is different if we are cued to pick up the book relative to if we simply look at the same book on the library shelf.

A possible relationship between action and episodic memory is suggested by action-related neuronal responses in two brain areas: the medial temporal lobe (MTL) and the locus coeruleus (LC). MTL structures, particularly hippocampus, are critical for episodic memory and spatial navigation^5,6^. In rodents, hippocampal theta rhythmic activity has long been associated with gross voluntary types of movement such as rearing and jumping^7^. Furthermore, the activity of hippocampal place cells, which fire when the animal visits a specific area in a familiar environment^5^, is also strongly dependent on movement-related information^8^. In humans, intracranial recordings from the medial temporal lobe reveal that voluntary movements of the arm or tongue, in contexts not requiring explicit memory encoding, modulate neuronal firing rates in hippocampus^9^ and surrounding cortex including parahippocampal gyrus^10^. These examples indicate that movement modulates neural activity in the MTL, a region critical for episodic memory, suggesting that action may affect memory formation.

The LC is the brain’s main source of noradrenaline (NE), a neuromodulator known to modulate episodic memory^11–14^. Single-unit recordings of the LC in non-human primates^15–18^ and cats^19^ demonstrate increased activity with goal-directed actions. This raises a possibility that the NE released by action-induced LC activity may promote on-going cognitive functions, such as the encoding into episodic memory of stimuli presented simultaneously with the action. We therefore hypothesized that taking action would enhance episodic memory encoding by engaging medial temporal lobe memory circuits via recruitment of the noradrenergic system.

To test this hypothesis, we examined how encoding of a visual stimulus is influenced by simultaneous voluntary movement (“Go” button-press response) compared to withholding of movement (“NoGo” response). A total of 237 healthy, young participants were tested over a series of experiments, that included functional magnetic resonance imaging (fMRI) and pupillometry studies, with different task manipulations (Table 1). The experiments started with an encoding task during which participants viewed pictures with the requirement to perform an action (Go-items) or withhold an action (NoGo-items) indicated by the color of a surrounding frame. Participants subsequently performed a surprise recognition task (Figure 1). Initial behavioral experiments confirmed our prediction that action modulates memory encoding. Subsequent experiments employing fMRI, pupillometry and manipulation of the emotional content of encoded material provide converging evidence to support our hypothesis that Go-associated encoding enhancement is mediated by the adrenergic system.

**Figure 1.**
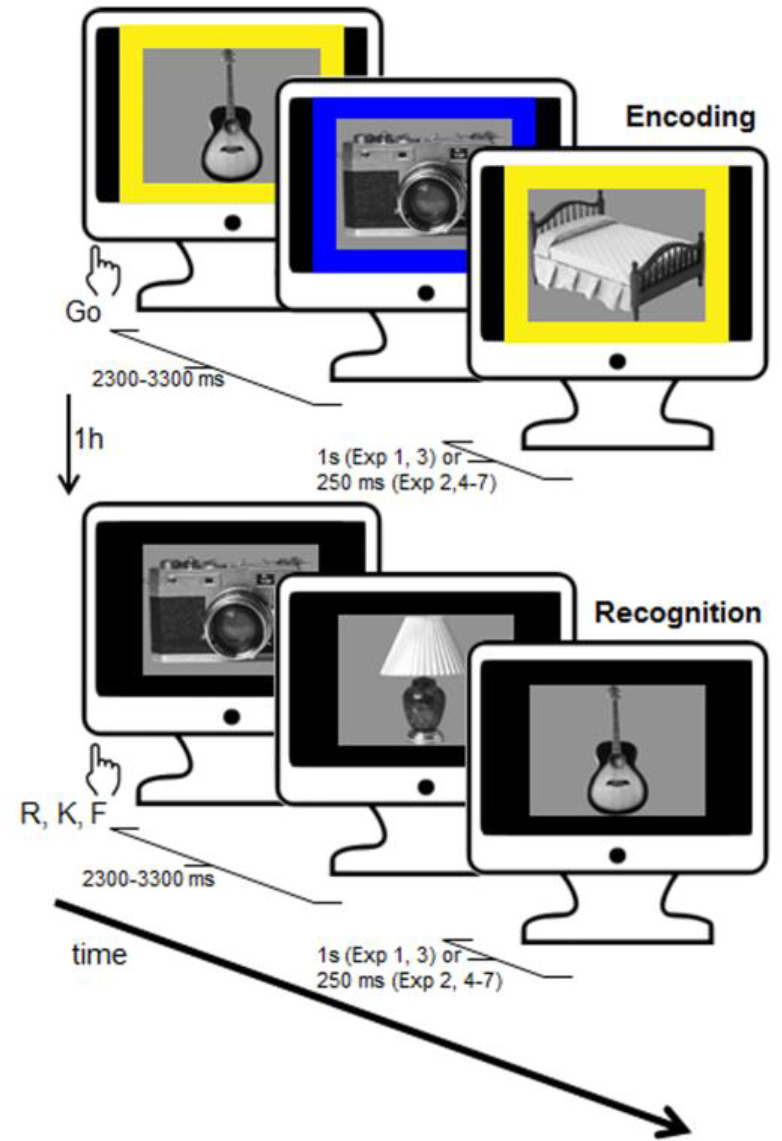
Behavioral task: Incidental memory encoding in the context of a Go/ NoGo task. At encoding, grayscale objects were presented with a color fame indicating requirement of a button press for the Go condition or withholding the response for NoGo. A surprise recognition test was conducted one hour later (or one day later Exp 7), during which participants were presented with objects from the encoding task intermixed with an equal number of lure items (presented without a frame) and indicated whether they remembered (R), were familiar with (K) or did not remember (F) the objects.

**Table 1.** Summary of experimental protocol, participants and experimental context for Exp 1-7. Presentation time pertains to both encoding and recognition tasks. *N*: Number of participants.

| Exp | Presentation time | Recognition test interval | Performance financially rewarded | Context | *N* | Emotional stimuli |
| --- | --- | --- | --- | --- | --- | --- |
| 1 | 1s | 1h | no | behavior | 31 | No |
| 2 | 250 ms<br>(0/250/500ms) | 1h | no | behavior | 38 | No |
| 3 | 1s | 1h | yes | behavior | 26 | No |
| 4 | 250 ms | 1h | no | behavior | 22 | No |
| 5 | 250 ms | 1h | no | fMRI | 21 | No |
| 6 | 250 ms | 1h | no | pupillometry | 28 | No |
| 7 | 250 ms | 1 day | no | behavior | 21 | Yes |

## RESULTS

### Taking action boosts episodic memory

In the first experiment (Exp 1), we tested for the effect of performing an action during encoding on subsequent memory performance. During the surprise recognition test, participants were required to make “remember”, “know” or “new” (R/K/N) judgments^20^, with remember responses indicating recall of elements of the study episode, know responses indicating a sense of familiarity, and new responses indicating the picture was not presented at encoding. We observed significantly better recollection for items requiring Go responses at encoding compared to NoGo-items (*t*_30_ = 2.40; *P* = 0.023), (Table 2). Successful encoding of Go items was not modulated by response speed, as reaction times (RTs) for subsequently remembered and forgotten Go items did not differ (Figure 2b, Supplementary Table 1). By contrast, memory performance for familiarity judgments was at chance level for both Go and NoGo stimuli (one-sample *t*-tests K hits minus false alarms, *P*_S_ > 0.49) and did not differ between them (*t*_30_ = 0.55; *P* = 0.587). This absence of a relevant memory signal for K responses, although contradicting previous literature^21^, was generally the case for all subsequent experiments (Supplementary Table 2, Supplementary Figure 1). This possibly reflects the difficulty of the memory task given that participants could be focusing more on the cue frame than on the picture. We therefore focused all further analyses on remember accuracy and its modulation by motor response at encoding. We replicate action-induced enhancement of remember accuracy across 6 subsequent variants of this experiment (Figure 2a-b, Table 2).

**Figure 2.**
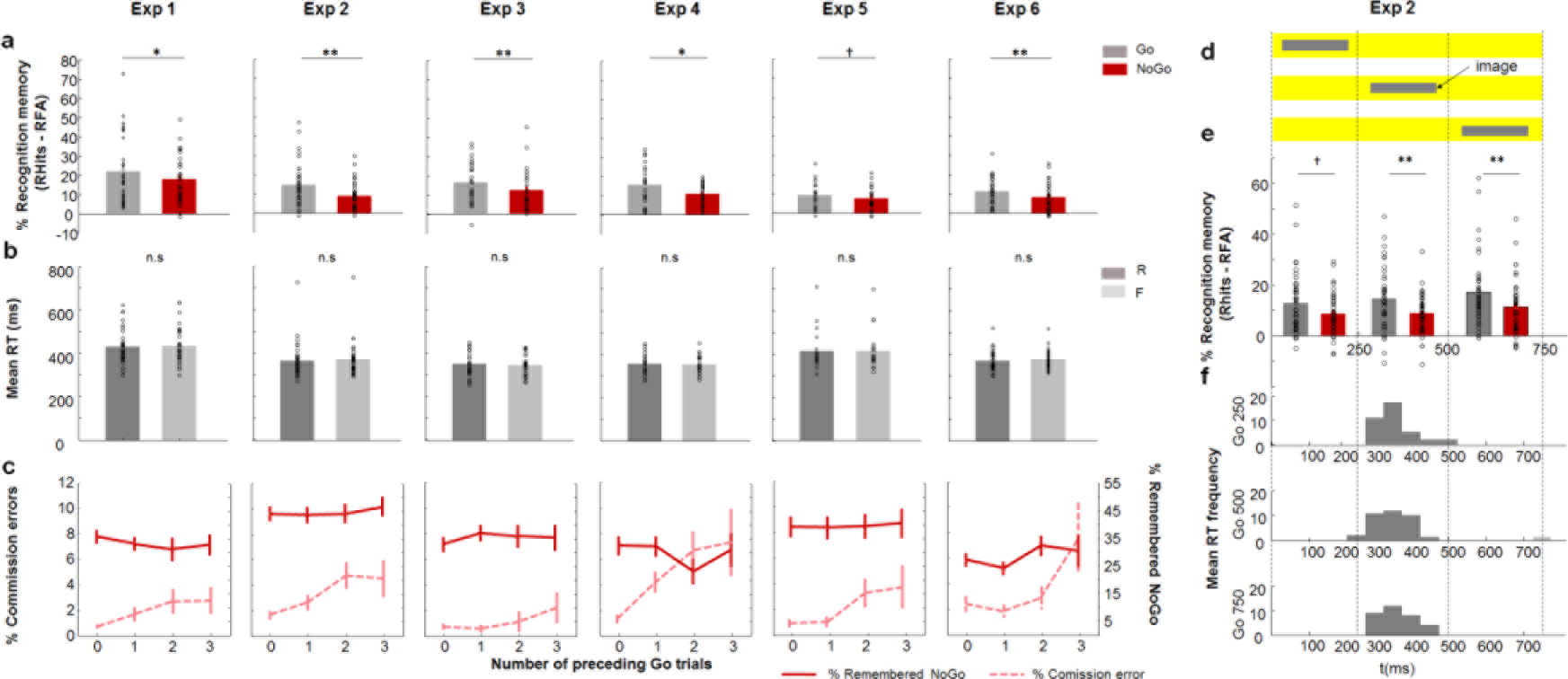
(See also Supplementary Figure 1, Supplementary Tables 1 and 3). Behavioral results: Exp 1-6 (a) Memory performance. Recognition memory for remembered (R) items corrected by false alarms (proportion of remembered (R) responses to new items) for both Go and NoGo conditions for each experiment * (*P* < 0.05), ** (*P* < 0.01), † (*P* < 0.05 one-tailed). (b) Reaction times at encoding for remembered and forgotten Go stimuli. (c) Commission error rate on NoGo trials during encoding depending on the number of consecutive preceding Go trials in black and Recognition memory (remember hits minus false alarms) for the NoGo condition depending on the number of consecutive preceding Go trials in red. (d) Experimental design of Exp 2. The colored background indicating Go or NoGo response requirement was presented for 750 ms while grayscale objects were presented for 250 ms on the center of the screen in one of 3 possible onset times: 0, 250 or 500 ms. (e) Recognition memory for remembered items corrected by false alarms for each condition (Go/NoGo) and the time window of presentation (250/500/750ms). (f) Histograms for the mean RTs across participants for Go condition at each time window of presentation.

**Table 2.** Summary of paired *t*-test results comparing remember accuracy (% remembered items minus false alarm rate) for Go vs. NoGo stimuli for Exp 1-6.

| **Exp** | ***t*-test Go vs. NoGo** | ***P* value** | **Cohen's *d* Go vs. NoGo** |
| --- | --- | --- | --- |
| 1 | $t_{30} = 2.40$ | 0.023 | 0.279 |
| 2 | $t_{37} = 3.28$ | 0.002 | 0.566 |
| 3 | $t_{25} = 2.85$ | 0.009 | 0.373 |
| 4 | $t_{21} = 2.26$ | 0.034 | 0.509 |
| 5 | $t_{20} = 1.41$ | 0.175 | 0.293 |
| 6 | $t_{27} = 2.75$ | 0.010 | 0.397 |

### Taking action enhances episodic memory, inhibiting action does not impair it

Memory was better for Go- compared to NoGo-items suggesting that taking action enhances encoding. However, as memory performance is compared between two response requirements at encoding (Go *vs*. NoGo), the difference in memory could alternatively be explained by response inhibition resulting in memory impairment. To test this alternative account, we utilized the fact that, due to the cue to Go or not being randomized across trials, the number of consecutive Go trials preceding NoGo trials was variable. With increasing consecutive Go trials, response inhibition mechanisms are more taxed, leading to increased commission errors (*i.e.*, a “Go” response when “NoGo” is cued)^22,23^. By extension, an inhibitory mechanism underlying NoGo-evoked worsening of memory encoding would predict that memory for NoGo items would decrease with increasing preceding number of Go items. We did not observe this relationship. Although participants in Exp 1 indeed showed more commission errors as a function of the number of consecutive preceding Go trials, showing a linear increase (*F*_1,30_= 5.44; *P* = 0.027; partial *η*^2^ = 0.349) (Figure 2c; Supplementary Table 3), no effect of the number of preceding Go trials on NoGo-item memory was found (*F*_2.34,70.16_ = 0.82; *P* = 0.487). This lack of NoGo-item memory modulation on the basis of preceding Go trials, also observed in all subsequent experiments (Figure 2c; Supplementary Table 3), argues against an inhibitory mechanism negatively affecting memory for NoGo items.

### Action-induced memory enhancement is not dependent on stimulus presentation time window

To provide further evidence that the mnemonic difference in memory performance between Go and NoGo-associated stimuli results from an action-induced memory enhancement and not from a NoGo-induced (action inhibition) encoding impairment, Exp 2 tested if temporal overlap between stimulus presentation and putative inhibitory neural responses would determine memory performance. Critically, inhibition during Go/NoGo tasks is linked to changes in the amplitude and topography of different waveforms of event-related potentials (ERPs) peaking at ~200-300 ms^24,25^. Exp 2 therefore employed a variable temporal asynchrony between cue-frame and grayscale picture presentation. In this experiment, grayscale pictures for both conditions (Go and NoGo) were presented during one of 3 consecutive time windows of 250 ms (0-250, 250-500, 500-750 ms; Figure 2d) with the frame-cue presented from 0-750 ms. Importantly, an inhibitory account of NoGo-induced encoding disruption predicts that poorer NoGo than Go memory would be most pronounced in the earliest stimulus presentation window (0–250ms), as this corresponds to the temporal profile of inhibition response-associated electrophysiological activity (*e.g.*, the N2 component).

Directly contradicting this account, Go item memory was better than NoGo item memory at all stimulus presentation windows (Figure 2e), with similar reaction time (RT) distributions for the 3 windows (Figure 2f). In a first analysis, collapsing over presentation time window, we replicate a significant effect of motor response on subsequent remember accuracy (*t*_37_ = 3.277; *P* = 0.002, Figure 2a). Critically, a repeated measures ANOVA with factors memory performance (remembered Go, NoGo) and time window of presentation (0, 250 or 500ms) showed no significant interaction (F_24.615,73.334_ = 0.22; *P* = 0.802). This suggests that the effect of voluntary movement/withholding movement on subsequent memory does not occur exclusively at a particularly early stage of the inhibitory process (Figure 2e). We note that Go-associated memory is greater than NoGo memory even when stimuli are presented 500 ms after cue frame onset, which is later than all RTs for this stimulus type (Figure 2f). Although other NoGo-related ERPs occurring later than the N2 have been described, the results of Exp 2 together with the observed lack of NoGo-item memory modulation on the basis of preceding Go trials, provide evidence against inhibition-induced memory impairment.

### Taking action enhances episodic memory regardless of reward anticipation

Human neuroimaging data suggest that memory formation is promoted by anticipation of reward through interactions between MTL structures and dopaminergic midbrain^26,27^. Furthermore, there is recent evidence that button-press Go responses in anticipation of reward can improve memory encoding^28^. This raises a possibility that Go responses in the current task reflect an approach-related action that engages the reward system which, in turn, strengthens episodic memory. If this were the case, the ensuing prediction is that explicit anticipation of financial reward would evoke greater action-evoked memory enhancement than observed in Exp 1. Thus, the design of Exp 3 was identical to Exp 1 (frame-cue and picture presented simultaneously) except that participants were financially rewarded for responding as fast as they could, and financially penalized for omission and commission errors. These task instructions led to significantly faster RTs for Go trials in Exp 3 than in Exp 1 (*t*_55_ = 4.57, *P* < 0.001). Again we demonstrate significantly better remember accuracy for Go– *vs.* NoGo-items (t_25_ = 2.85; *P* = 0.009; Figure 2a). However, a repeated measures ANOVA on remember accuracy with within-subjects factor response type (Go, NoGo) and between-subjects factor experiment (Exp1 No Reward, Exp 3 Reward) did not show a significant interaction (*F*_1,55_ = 0.01; *P* = 0.913). The main effect of Go *vs.* NoGo memory was significant (*F*_1,55_= 12.65; *P* = 0.001; η^2^ = 0.187), with effect sizes for the Go vs. NoGo memory comparisons comparable across the two tasks (Table 2). There was not a significant between-subject effect of Exp (*F*_1,55_= 2.65; *P* = 0.109). These results therefore indicate that there is no additive effect of reward anticipation on the observed action-induced memory enhancement.

### Action-induced episodic memory enhancement is associated with increased Locus Coeruleus BOLD activity

Our behavioral studies were predicated on the hypothesis that taking action would boost episodic memory via interactions between the noradrenergic system and medial temporal lobe memory circuits. To test this mechanistic hypothesis, we conducted a functional magnetic resonance imaging (fMRI) study. First, a behavioral pre-fMRI experiment (Exp 4) was performed, identical to Exp 1 but with presentation duration of 250ms. Exp 4 simply ensured that robust memory enhancement for Go *vs*. NoGo stimuli is observed at this shorter presentation time, employed so as to minimize saccades, which not only lead to spurious BOLD effects^29^ but also represent another type of action that increases LC activity in non-human primates^17^. The results of Exp 4 once more replicated the main finding from Exp 1-3 of better memory for Go *vs*. NoGo items (*t*_21_ = 2.26; *P* = 0.034; Figure 2b).

The task employed in the context of fMRI scanning (Exp 5) was identical to Exp 4. Behaviorally, although there was a remember advantage for Go *vs*. NoGo stimuli, this effect was not significant (*t*_20_ = 1.41; *P* = 0.175). The primary aim of this fMRI experiment was to derive mechanistic insights into memory enhancement for stimuli paired with action. Testing for an interaction between motor response (Go *vs*. NoGo) and subsequent memory (R *vs*. F) identified a significant activation in dorsal pons, in an area consistent with LC (Figure 3a, Supplementary Table 4). Note that this effect was also observed if the sample was restricted to the 14 subjects showing go-induced memory enhancement. To increase the robustness of spatial localization of this response to LC, we repeated this analysis using an infra-tentorial template for spatially unbiased, nonlinear normalization of brainstem and cerebellum (SUIT) to provide more accurate intersubject-alignment of the brainstem than whole-brain methods. A significant action by subsequent memory interaction was again observed in dorsal pons, overlapping with a probabilistic atlas of the LC (Figure 3b).

**Figure 3.**
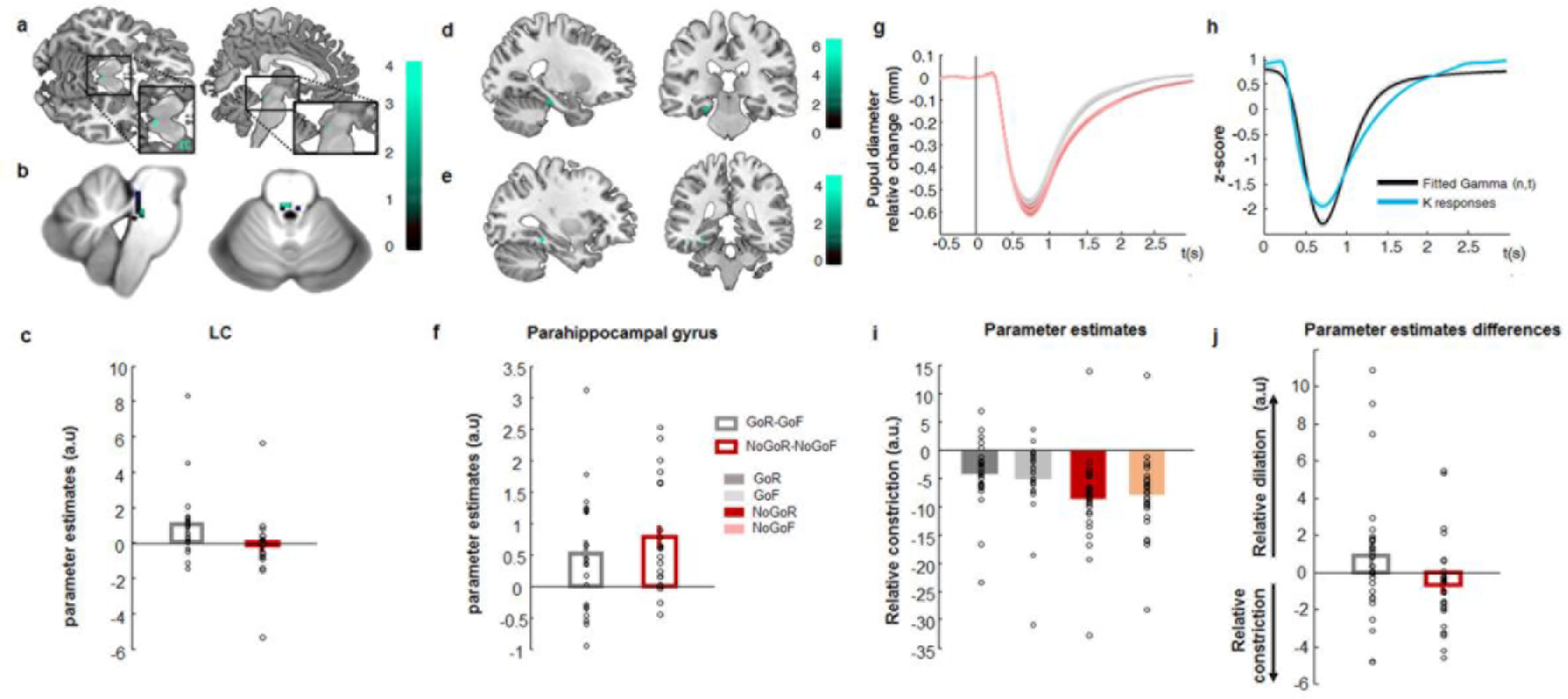
(See also Supplementary Figure 2, Supplementary Tables 4, 7 and 8) Go responses during successful encoding engage the noradrenergic system. (A-C) LC activation (Exp5). (a) Interaction between action (Go vs NoGo) and subsequent memory (remembered vs forgotten) has been overlaid on a canonical T1 image (threshold for illustration, here and in subsequent panels, *P* < 0.001 uncorrected) to show activation in an area of the dorsal pons consistent with LC (2 −28 −16; Z = 3.38; *P* < 0.001 uncorrected). (b) The same interaction, normalized to the SUIT atlas template (−2, −34, −23, Z = 3.32; SVC *P*_FWE_ = 0.035) is overlaid on a high resolution atlas template of the human brainstem and cerebellum, with a probabilistic spatial mask (at 1 std) of LC superimposed. (c) Parameter estimates for the BOLD response in LC. Error bars pertain to s.e.m. (d-f) Parahippocampal activation (Exp5). (d) The comparison between remembered vs. forgotten items reveals parahippocampal activation (−24 −26 −20; Z = 3.73; SVC *P*_FWE_ = 0.05), shown overlaid on a canonical T1 image. (e) A psychophysiological interaction between LC activation and action-induced modulation of memory is significant in parahippocampal gyrus (−32, −38, −12; Z = 3.79; SVC *P*_FWE_ = 0.02). (f) Parameter estimates for the parahippocampal activation shown in D. (g-j) Pupillary responses during successful encoding are modulated by Go responses (Exp 6). (g) Raw pupil diameter relative to baseline change measures averaged over participants (shaded error bars pertain to s.e.m.). (h) The Erlang gamma function for the early visual component (black) parameterized by fitting to the z-scored pupil responses at encoding to subsequent familiar (K) responses (blue). (i) Parameter estimates for the cognitive aspect of the pupil responses for GoR, GoF, NoGoR, NoGoF conditions (error bars pertain to residual error of the model). (j) Parameter estimates for the differences of the relative pupil dilation/constriction for the interaction between motor response (Go, NoGo) and subsequent memory (R, F) for the cognitive aspect of the pupil dilation.

Furthermore, both the Go *vs*. NoGo main effect comparison (Supplementary Figure 2a, Supplementary Table 5) and the opposite test (Supplementary Figure 2b, Supplementary Table 6), revealed cortico-subcortical networks previously shown to be involved in action and response inhibition^30,31^, respectively. As predicted, MTL activation, in parahippocampal gyrus, was observed in the R *vs*. F, subsequent-memory comparison (Figure 3d and f, Supplementary Table 7) as has been reported previously^32,33^.

LC sends widespread noradrenergic projections to cortical and subcortical structures, including the MTL (for review see^34^). To determine which regions correlate with LC activity during encoding, we performed a psychophysiological interaction (PPI) analysis to estimate context-specific changes in correlation between the LC and the rest of the brain. Specifically, we tested which regions were functionally connected with LC under the experimental context of successful encoding between Go *vs*. NoGo trials. Connectivity between LC activity and parahippocampal gyrus was observed (Figure 3e, Supplementary Table 8), in a region in close proximity with parahippocampal cortex expressing a main effect of successful object encoding (Figure 3d). These results suggest that action-evoked memory enhancement is mediated by a LC-parahippocampal gyrus circuit: NE neuronal activity is triggered by action, as has been previously shown in animal studies^15–19^, and the ensuing NE release targets the MTL promoting memory formation^11–14,35^.

### Action-induced episodic memory enhancement is associated with increased pupil dilation responses

Our fMRI results indicate that action-induced memory enhancement results from noradrenergic LC responses that upregulate episodic memory encoding processes in the MTL. To provide a second, independent index of LC activity during Go-induced encoding enhancement, we next performed the same Go/NoGo memory paradigm while recording pupil diameter responses (Exp 6). Pupil diameter has been shown to be a reliable, indirect index of LC activation^36^, positively correlating with LC firing rates in non-human primates^37,38^, and with BOLD activity in human LC^39^. Furthermore, pupil diameter is also modulated by learning and memory processes^35,40^. We therefore expected that the pupil-derived index of LC activation would relate to memory performance and action in the same way as that observed with fMRI.

The behavioral results of Exp 6 again show better remember accuracy for stimuli paired with Go *vs.* NoGo responses (*t*_27_ = 2.75, *P* = 0.010) (Figure 2a, Table 2). Note that the behavioral task used in Exp 4, 5 and 6 was identical, with Exp 4 and 6 showing significantly better remember accuracy for Go *vs*. NoGo stimuli. To test for an interaction between encoding and button press in pupil diameter, the raw pupil responses (Figure 3g) were submitted to a general linear model (GLM) using a basis function approach. Stimulus-locked pupillary responses were modeled with two basis functions, one pertaining to the light-reflex and a later cognitive component. An Erlang gamma function was used for the later cognitive component. The earlier visual component was parameterized by fitting the pupil responses at encoding to stimuli later judged as familiar (K), as statistical inference was not made on these K trials (Figure 3h). For each subject, parameter estimates for regressors convolved with the cognitive basis function pertaining to our conditions of interest (GoR, GoF, NoGoR and NoGoF) were entered into a 2 by 2 ANOVA. Go responses evoked greater pupil dilation (main effect of Go *vs.* NoGo, *F*_1,27_ = 17.78; *P* < 0.001) as is clear from the raw pupil traces, where an initial pupil constriction, due to the light-reflex to stimulus presentation, is followed by a later differential dilation (Figure 3g), in line with previous observations in human^41,42^ and non-human primates^18^. Critically, a significant interaction between response type (Go *vs.* NoGo) and subsequent memory (R *vs*. F) is observed (*F*_1,27_ = 4.21; *P* = 0.050). As predicted from the LC activation in Exp 5, this interaction reflects greater pupil dilation to stimuli paired with Go responses that are subsequently remembered (Figure 3 i-j). These results, derived from using pupil diameter as a proxy measure of LC activation, reinforce our fMRI evidence that Go-induced encoding enhancement is mediated by the noradrenergic system.

### Action-induced episodic memory enhancement depends on arousal

Across all experiments we observed, at a group level of statistical inference, a consistent memory advantage for stimuli requiring a Go response at encoding. However, the advantage is not observed in all subjects. We hypothesized that this could be due to intersubject differences in arousal levels during performance of the cognitive task. This hypothesis is based on the inverted-U relationship between arousal (and noradrenergic activity) and cognitive performance (the Yerkes-Dodson law^43^). That is, if participants already show a certain degree of arousal during encoding, further Go-evoked NE release would be detrimental to encoding performance. To test this hypothesis, we performed an additional analysis on data recorded in Exp 6 to examine an index of arousal derived from pupil measures. The light-reflex (pupil constriction in response to light) shows reduced amplitude in patients with generalized anxiety^44^, and is reduced in healthy individuals in the context of arousal produced by pain expectation, with this decreased light reflex response correlating with increased subjective anxiety^45^. Our prediction was, therefore, that participants with highest level of arousal during task performance (*i.e*., those with reduced light reflex) would not show action-induced memory enhancement. To confirm this prediction, we calculated the light-reflex amplitude for all participants in Exp 6 by averaging across all encoding trials for each subject. Figure 4 shows the pupil response as an average function for two groups: the 22 participants completing Exp 6 who showed enhanced memory for Go vs. NoGo stimuli, and the remaining 6 participants who did not. There is a reduction in light-reflex amplitude in the participants not showing Go-induced memory advantage compared to the participants who do show a Go-induced memory advantage (*t*_10.747_ = −2.23; *P* = 0.048), supporting our explanation that these individuals have a higher level of arousal during encoding, which negates the mnemonic benefit afforded by Go-induced LC activity.

**Figure 4.**
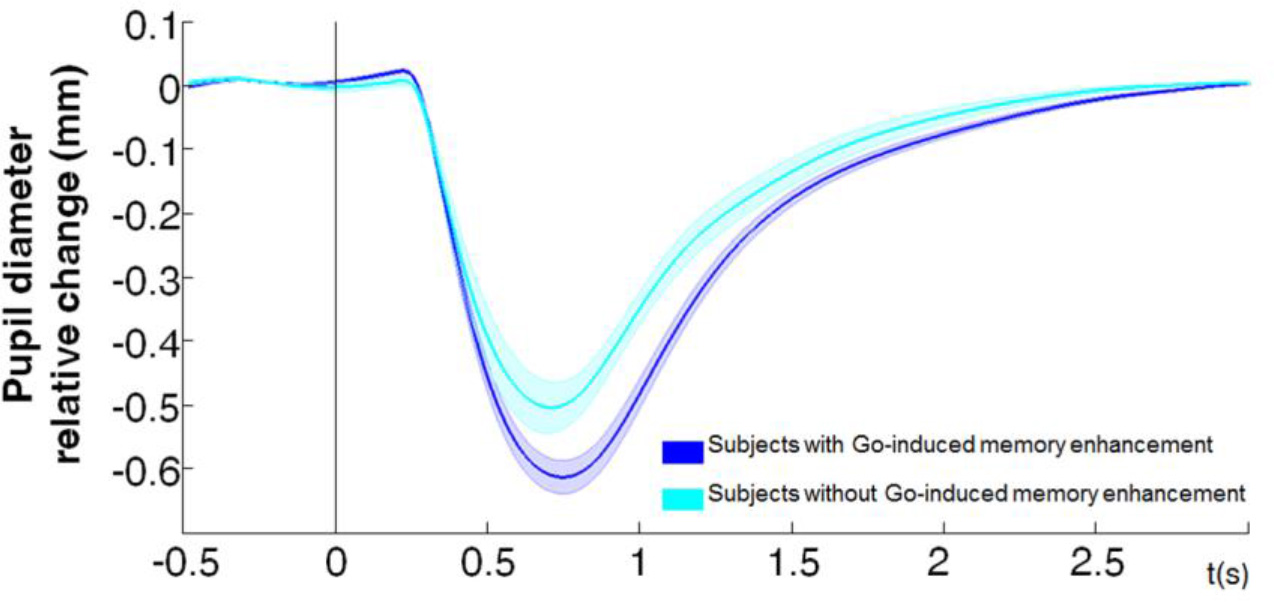
Reduced pupillary light-reflex constriction for participants that do not show Go-induced memory enhancement. Average pupil diameter change relative to baseline is plotted for two groups: the 22 participants completing Exp 6 who show enhanced memory for Go vs. NoGo stimuli (blue), and the remaining 6 participants who do not (light blue).

### Action-induced episodic memory enhancement is modulated by emotion

The results of Exp 5 and 6 demonstrate involvement of LC in Go-induced memory enhancement, indicative of an underlying noradrenergic mechanism. A NE mechanism has also been shown to underlie enhanced memory for emotionally negative relative to neutral stimuli^12–14,46^. Thus, if Go- and negative emotion-induced memory enhancements are both mediated by a NE mechanism, a novel hypothesis can be derived stating that Go-induced memory enhancement is modulated by the emotional nature of the stimulus presented simultaneously with movement. This rationale is again based on the inverted-U relationship between arousal (and noradrenergic activity) and cognitive performance^43^. That is, if during memory encoding there are two psychological parameters that increase noradrenergic drive (emotion and voluntary movement), these effects may influence encoding performance in a predictable way (Figure 5a). As emotional arousal increases, encoding performance for emotional stimuli (Go emotional and NoGo emotional) should move rightwards on the inverted-U curve beyond optimal NE effect on encoding. Highest noradrenergic drive, evoked by the condition involving both action and aversive emotion simultaneously (*i.e*., Go emotional) should thus worsen memory performance. Neutral stimuli (NoGo neutral, Go neutral) should lie on the left side of the curve, with lowest noradrenergic drive for the condition not involving action or aversive emotion (NoGo Neutral) showing lowest memory performance. By contrast, optimal memory performance should be situated in the middle of the curve corresponding with moderate level of emotional arousal and NE release driven by only one NE mediator, *i.e*., either action or emotion (Go Neutral, NoGo Emotional) (Figure 5a). The alternative hypothesis would be a simple summation of effects of NE drive, producing a main effect on both action and emotion, but no interaction.

**Figure 5.**
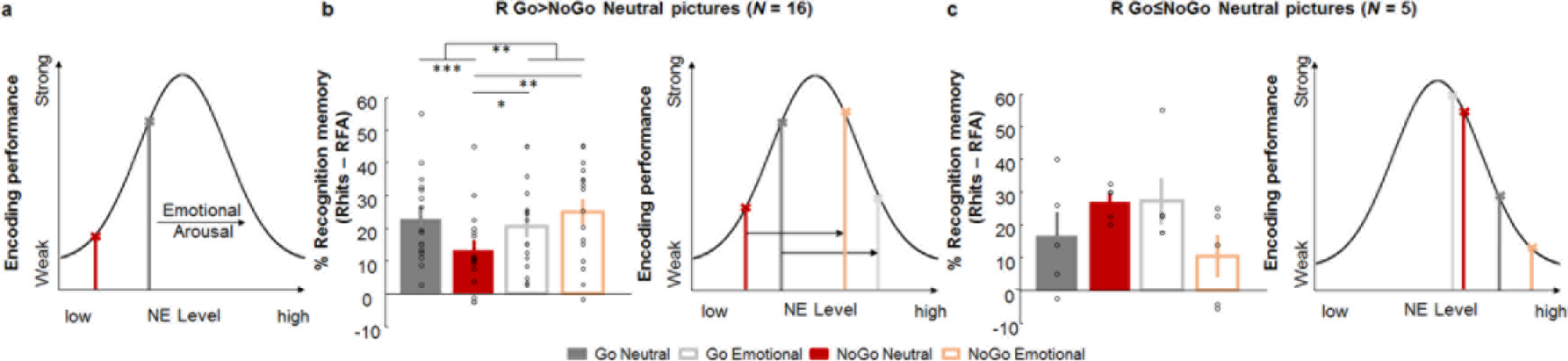
Go-induced encoding enhancement is modulated by emotion. (a) Schematic “inverted-U” relationship between encoding performance and norepinephrine (NE) level, with putative locus of Go and NoGo encoding for emotionally neutral stimuli indicated on this curve. We hypothesized that emotion would shift memory scores to the right. (b-c) Recognition memory for remembered items (R) corrected for false alarm rates for Go and NoGo neutral and emotional trials (left) and the schematic (right) for participants that show Go-induced memory enhancement for the neutral stimuli (*N* = 16) (b) and those that do not (*N* = 5) (c) * (*P* < 0.05), ** (*P* < 0.01), *** (*P* < 0.001).

To test these predictions, we performed a final experiment (Exp 7), which was identical to Exp 4 except that, instead of grayscale pictures of objects, participants were presented with an equal number of neutral and emotional color scenes from a standardized database. The cue to Go or NoGo was again indicated by a blue/yellow frame. The enhancing effect of emotion is known to be greater when memory is tested after long (considered to be from 1h to 24h or more) than after short immediate intervals, thus the surprise recognition test was performed after a 24h delay to promote a greater effect of emotion on memory^47,48^. We first examined memory for participants showing Go-induced memory enhancement for neutral stimuli (16 of the 21 participants in total). This subgroup was selected in view of the results of Exp 6 showing that individuals not showing action-induced memory enhancement may already be at a heightened level of arousal, which could obscure additional memory effects of the emotional nature of stimuli presented at encoding. Strikingly, although this subgroup of participants show Go-induced encoding enhancement for neutral stimuli, this is not observed for emotional stimuli (Figure 5b). The Go-induced decrease in encoding of emotional pictures is in keeping with our predictions that the combination of emotion and Go-responses moved LC activity beyond the optimum of the inverted-U function for memory encoding (Figure 5b). An emotion (neutral, aversive) by response (Go, NoGo) ANOVA on encoding performance showed a significant interaction (*F*_1,15_ = 14.12; *P* = 0.002; η^2^ = 0.486), whereas the main effect of response was not significant and that of emotion only showed a trend (*F*_1,15_ = 3.86; *P* =0.068). Post-hoc *t*-tests revealed significantly different memory performance between Go Emotional and NoGo Neutral stimuli (*t*_15_ = 2.88; *P* = 0.011) and NoGo Neutral and NoGo Emotional stimuli (*t*_15_ = −3.83; *P* = 0.002). The difference between Go *vs*. NoGo Neutral stimulus encoding (*t*_15_ = 7.34; *P* < 0.001) is obviously biased by preselection of participants showing this effect.

Interestingly, in Exp 7 those subjects that do not show Go-induced memory enhancement for neutral stimuli (*N* = 5), actually show better memory for NoGo neutral pictures (Figure 5c). If a Go-induced release of NE impairs memory in these participants, this would be compatible with these subjects operating more to the right of the inverted-U function of arousal. That is, they were in a state of higher arousal than other subjects during the course of the experiment (Figure 5c), in line with our findings from subjects in Exp 6 showing attenuated light-reflex. Again we find an opposite pattern for Go/NoGo effects on emotional stimuli (Figure 4c), although we predicted less memory for Go emotional that NoGo emotional in these subjects. Indeed, there was a trend towards a significant interaction between emotion and motor response (Friedman test *χ*^2^(3) = 7.41; *P* = 0.060). We note, however, the small sample size of participants showing this pattern. Nonetheless, overall the results of Exp 7 indeed confirm our predictions, based on the Yerkes-Dodson law, for memory performance showing an action-emotion interaction following an inverted-U for individuals with putatively normal levels of arousal. Moreover, they provide further support for a NE basis of action-induced memory enhancement.

## DISCUSSION

Over a series of experiments, we consistently observed better memory for stimuli cooccurring with action. Given that in all experiments, memory for Go stimuli was compared to NoGo items, we employed two strategies to make the case that the memory difference between these two stimulus classes reflected enhanced Go, and not impaired NoGo, encoding. First, we showed that increasing inhibitory load did not disrupt successful memory encoding, despite increasing commission error rates. This contradicts a recent suggestion of response inhibition-induced episodic memory impairment^49,50^. Indeed, this previous study showed that participants committed more NoGo errors with increasing number of preceding Go stimuli, but did not test for an expected increased disruption of memory as inhibitory load increased. We note that increasing inhibitory load with increasing consecutive Go trials increases the surprise elicited by the subsequent NoGo stimulus, and (working) memory can be impaired after surprising events that trigger motor inhibition^51^. The role of surprise, which is also associated with increased noradrenergic activity^34^, is however, small in our task, given the equiprobable presentation of Go and NoGo stimuli. Second, we provide evidence that the observed memory difference did not occur exclusively at an early stage of the inhibitory process corresponding to the temporal profile of response inhibition-evoked electrophysiological activity (*e.g*., the N2 component)^24,25^. Both strategies indicated that an inhibitory effect of action inhibition is unlikely to account for the encoding difference between Go and NoGo stimuli.

Data from fMRI and pupillometry experiments provided converging evidence for LC engagement as the mechanism underlying action-induced memory enhancement, implying a role for noradrenaline release in this process. Given the established role of noradrenaline in consolidation of memory for emotional experiences^11–14^ we introduced an emotional manipulation to our task (Exp 7), and showed that action-induced memory enhancement is also modulated by emotional arousal. Strikingly, the emotional content of stimuli interacted with putative action-driven NE release to modulate memory performance in a way that reflects Yerkes-Dodson law. This is in keeping with early studies showing that levels of arousal interact with injected NE dose to modulate memory performance following an inverted -U curve in rats^52^, meaning that low doses of NE do not alter, moderate enhance and high doses impair later retention performance. Together with our results that participants with higher levels of arousal (lower light-reflex amplitude) during task performance did not show action-induced memory enhancement, suggests that inter-subject memory variability can be explained by arousal state.

An LC-centered mechanism for Go-induced memory enhancement facilitates the interpretation of two further behavioral findings, when considered in the context of LC recordings in non-human primates. Reaction times did not differ between subsequently remembered and forgotten Go items, suggesting that the speed of response does not modulate action-induced memory enhancement. This absence of RT modulation mirrors findings in monkeys that the magnitude of LC firing during action is not modulated by RT^15–17^. Furthermore, in the current study, we found no additive memory benefit for financially rewarded, relative to unrewarded, Go trials. This is in line with monkey data showing LC firing aligned with action in the absence of reward anticipation; LC responses are observed with actions to no reward cues^16^, on non-rewarded trials^53^, with rewarded decision to act but not with rewarded decision to stop a response^17^. These observations argue in favor of the sufficiency of noradrenergic drive mediating the memory enhancement shown here.

Over a series of behavioral experiments, we provide the first empirical evidence that action performance can boost episodic memory for stimuli unrelated with the movement. By dissociating the action from the content of the memory, we provide a novel dimension to the enactment effect^3,4^. In Exp 2, stimuli were presented asynchronously with the cue for motor response, limiting the likelihood that action-induced memory enhancement simply reflects subjects remembering the association between an image and the action. Furthermore, the fact that button-presses did not influence when or how visual stimuli were presented differentiates our findings from the memory benefit observed during volitional control tasks, in which participant actions lead to self-controlled viewing, as opposed to passive viewing, of encoded stimuli^54^. Furthermore, the fact that subjects did not have to choose between two alternative button press responses differentiates our findings from previous results showing that the act of choosing enhances declarative memory^55^.

Our findings are supported by previous reports of enhanced memory for target-paired stimuli that require a button press^56–58^. This memory enhancement has been interpreted in terms of an attentional boost effect^59^. Given the critical role of the noradrenergic system^34^ for attentional processes, this interpretation can be accommodated by action-induced memory enhancement mediated by recruitment of the locus coeruleus. An explanation of Go vs NoGo memory advantage on the basis of Go responses being more attentionally demanding is unlikely, given that NoGo responses also require attention, particularly in the context of financial penalization of commission errors (Exp 3). It should also be noted that studies showing enhanced memory for target-paired stimuli typically employ target-detection tasks, where the stimuli requiring action are infrequent, thereby producing an “oddball” effect which is known to improve memory^60^.

Applying the mechanistic framework provided here to the everyday example given earlier, taking a book from the library shelf will trigger LC activity. The subsequent noradrenaline release will target parahippocampal gyrus to promote encoding and facilitate memory formation for the episode, including action-irrelevant aspects such as the title of the book. Thus, converging evidence presented in the current study argues for a relationship between LC and memory, proposing for the first time action as a link between them. These results provide a functional framework for potential future rehabilitation strategies using actions to enhance memory via noradrenergic engagement in individuals with memory impairment. Moreover, given the profound role of NE on cognition^34^, our observations likely extend beyond the memory domain and implicate action-induced modulation of a range of cognitive processes.

## AUTHOR CONTRIBUTIONS

Conceptualization, M. Y., M.K. and B.S.; Investigation, M. Y., A.G., J.G-R, V.S-L. and A. O.; Methodology, M.Y., B.S., A.O.de B, S.B. and M.K.; Software, M.Y.; Formal Analysis, M.Y. and B.S.; Writing – Original Draft M.Y. and B.S.; Writing – Review & Editing, B.S, J.G-R, M.K and S.B; Supervision and Funding Acquisition, B.S.

## ACKNOWLEDGMENTS

This work was supported by Project grant SAF2011-27766 from the Spanish Ministry of Science and Innovation and Marie Curie Career Integration Fellowship (FP7-PEOPLE-2011-CIG 304248) to B.S. M.K. is supported by an H2020 Marie Sklodowska-Curie fellowship and a Society in Science-Branco Weiss fellowship.

## ONLINE METHODS

### Participants

A total of 237 human subjects (aged 18-35; 116 female) were recruited via advertisement to participate in our study. Participants were right- or left-handed for the behavioral experiments and all right-handed for the fMRI experiment, had no history of neurological or psychiatric disease, and normal or corrected-to-normal visual acuity. All participants provided written informed consent prior to commencement of the study. The study was approved by the ethical committee of the Universidad Politecnica de Madrid.

### Psychological task

All experiments consisted of two phases: visual stimulus encoding in the context of a Go/NoGo task followed by a later surprise recognition test.

For Exp 1-6, from a pool of 380 grayscale photographs of objects, 190 were randomly selected and presented in randomized order during encoding. Participants were instructed to press a button (go trials) when the images were presented with a particular color frame (blue or yellow). The frame color for the “Go” and “NoGo” instruction was balanced across participants in all experiments. Go and NoGo frames appeared with equal probability (*i.e*., both at 50% probability). Participants were instructed to look at the center of the screen. Stimuli were displayed with 20 degrees of visual angle at a viewing distance of 60 cm. Participants returned after one hour in order to perform a surprise recognition test. For Exp 1-6, a total of 380 images – the 190 that were presented at encoding and 190 new “foils” – were presented in randomized order with no frames on a black background. Participants were required to indicate whether they remembered (R), were familiar with (K) or did not remember (forgotten, F) the image from the encoding phase (Figure 1).

For Exp 7, participants performed a similar Go/NoGo task but with color images selected from the International Affective Picture System (IAPS) Database^1^. We selected a total of 80 images, 40 neutral and 40 negative emotional, with the following arousal and valence ratings: emotional stimuli arousal score (std) 6.46 (0.49) and valence score 2.27 (0.85); neutral stimuli arousal score 2.89 (0.41) and valence score 5.00 (0.24), using a scale from 1-9 with 1 being the most negative valence and 9 most arousing. At encoding, participants were presented with 20 emotional and 20 neutral images (randomly selected from the pool of 80 images). Again, images were presented in random order with a color frame indicating the requirement of pressing a button for the Go trials or not pressing for the NoGo trials. A surprise recognition phase was conducted 24 hours after, instead of the 1h interval in all previous experiments, as the enhancing effect of emotion on memory is more evident at this longer time interval^2,3^. During recognition phase all 80 images were presented in random order. Again, participants made a Remember/Familiar/New judgment for each stimulus.

For all experiments, exclusion criteria were applied on the basis of task performance at encoding and recognition. Participants performing at less than 90% correct button press for Go, and 90% correct withheld responses for NoGo, trials were excluded from analyses. Furthermore, those participants with poor memory performance (defined as correct hit remembered rate minus remember false alarm rate less than 0%) were not further considered for analysis. In addition, participants making button-press responses for less than 90% of trials during recognition testing were excluded from analyses.

### Exp 1

Image presentation time was 1s in both phases, with variable ISI from 2.3 to 3.3s at encoding and 2.8 to 3.3s at recognition. A total of 33 participants (18 women; 32 right handed; age range, 18-35 years; mean age, 26.80 years; SD, 3.20) performed the experiment. Two participants were excluded on the basis of Go/NoGo task performance.

### Exp2

In this experiment, the image presentation time was reduced from 1s to 250ms. While a colored background indicating the requirement of pressing or not pressing a button was presented for 750 ms, the images of grayscale objects were presented at 3 different onsets relative to the 0 to 750 ms color background presentation time: at 0s, at 250ms and at 500ms. The same ISI was used as in the previous experiments. Up to 52 participants performed the experiment and 38 participants (22 females; 35 right handed; age range, 18-35 years; mean age, 27.97; SD, 4.26) were included in the final analysis. Five participants were rejected for poor Go/NoGo performance, 2 participants were rejected because of multiple button presses for the Go trials, 1 did not finish the task, 2 participants misunderstood the instructions and pressed when the image appeared not when the colored frame appeared on the screen, and 4 more were excluded on the basis of poor memory performance.

### Exp 3

This was identical to Exp 1 except that participants were financially rewarded for responding as fast as they could, financially penalized for omission and commission errors. Participants began the experiment with a 10€ voucher and were told they could earn up to 20€ on the basis of their performance. They were informed that at the end of the experiment, 5 Go trials and 5 NoGo trials would be randomly selected and for each correct NoGo, 1€ will be added to the final quantity, 0€ otherwise. For each correct Go trial they started with 1 more euro and per 0.1s of delay above 0.5s, 0.1€ were subtracted for the final amount. 29 participants (14 females; 28 right handed; age range, 18-35 years; mean age, 24.03; SD, 4.19) performed this experiment. One participant was discarded due to absent button-presses at recognition, and a further 2 participants for poor memory performance. All participants were paid 20€.

### Exp 4

This was identical to Exp 1 except that stimuli were presented for 250ms. 27 participants (16 females; 26 right handed; age range, 18-35 years; mean age, 26.74; SD, 4.92) took part in this experiment. Five participants were removed from the analysis: 3 on the basis of Go/NoGo performance and 2 were excluded on the basis of low response rate during recognition.

### Exp 5

This experiment is a replication of Exp 4 but in the context of fMRI scanning. Thirty-eight participants (25 females; 38 right handed; age range, 18-35 years; mean age, 25.34; SD, 5.06) began this experiment. On the basis of stimulus-correlated head movement we excluded 6 participants at encoding phase, 1 other subject was excluded due to signal drop out in medial temporal lobe, 6 other participants were rejected because they presented less than 85% of correct Go or NoGo trials or answers at recognition phase, 1 other because multiple button press for the Go condition and 3 were removed because of poor memory performance.

fMRI data acquisition. For each subject, a 3T Siemens Trio TIM system was used to acquire MPRAGE T1-weighted anatomical images with 1mm^3^ resolution (repetition time (TR), 2300 ms; echo time (TE), 2.98 ms; flip angle, 9°). During encoding, 288 gradient-echo echo-planar T2*-weighted MRI image volumes with blood oxygenation level-dependent contrast were acquired, plus five additional volumes, acquired at the start of each session and subsequently discarded, to allow for T1 equilibration effects. Each whole-brain volume comprised 40 axial slices (2.2mm thick; distance factor 0.25; repetition time 2.43 s; echo time 30 ms; flip angle 90°, FOV 192 mm × 192 mm; matrix 64 × 64) sequentially acquired (ascending).

fMRI data analysis. Functional imaging data were analyzed using statistical parametric mapping (SPM8; http://www.fil.ion.ucl.ac.uk/spm) using an event-related design. Each subject’s fMRI time series was realigned, slice time corrected, normalized to MNI space and smoothed with an isotropic 3D Gaussian kernel of 6 mm full-width half-maximum. To test for effects of motor action on memory, we specified 6 effects of interest in a general linear model (GLM): the events corresponding to Go and NoGo trials, separated according to whether these images yielded a subsequent remember (R), familiar with (K) or forgotten (F) response at recognition testing. Event-specific responses were modeled by convolving a delta-function with a canonical haemodynamic response function (HRF) to create regressors of interest. Response errors were modeled separately. Six movement parameters were modeled as nuisance covariates.

Session-specific parameter estimates of the magnitude of the hemodynamic response for each stimulus type were calculated for each voxel in the brain. A contrast of parameter estimates modeling each comparison of interest (e.g., remembered *vs*. forgotten Go *vs*. NoGo images) was calculated in a voxel-wise manner to produce, for each subject, one contrast image for that particular effect. For the random effects analysis, each subject's contrast image was entered into a one-sample *t*-test across participants. We report group- level analyses pertaining to the main effects and interaction term of our response (Go, NoGo) by subsequent memory (Remembered, Forgotten) 2 by 2 factorial design.

In order to improve the spatial reliability of the observed LC response, the SUIT toolbox was employed, as described previously^4,5^. In brief, realigned, slice-time corrected functional images were coregistered to their subject-specific T1-weighted anatomical scan (with origin manually set at the anterior commissure). We then repeated the first level analysis described above. Again, a contrast image for the interaction term of response (Go, NoGo) by subsequent memory (Remembered, Forgotten) was calculated in a voxel-wise manner. Next, the cerebellum and brainstem were isolated in the anatomical image, and the ensuing image normalized to the SUIT atlas template using a nonlinear deformation. This deformation was then applied to the contrast image created for the interaction term, and resliced, masking out activation from outside the cerebellum or brainstem. Finally, each participant’s normalized contrast images were smoothed with a 6 mm kernel and submitted to a second level GLM across subjects.

We carried out a small-volume correction (SVC) to the *P* values of the ensuing maxima in LC and parahippocampal gyrus. For the latter, we used bilateral posterior parahippocampal mask from the Harvard-Oxford atlas in view of previous evidence for encoding-related responses in this area to pictures yielding subsequent high-confidence remember judgements^6^. For LC maxima, we used a probabilistic LC atlas^7^ normalized to the anatomical space define by the SUIT toolbox.

### Exp 6

This experiment was performed in the context of pupillary recordings. The behavioral task was identical to Exp 4, except for the presentation of a white fixation cross in the center of the screen for the last 500 ms of the trial to allow participants to blink. Thirty-five participants (21 females; 32 right handed; age range, 18-35 years; mean age, 27.31; SD, 5.08) performed the experiment. Three participants were excluded from analyses due to excessive number of blinks (more than 10% of trials in any one of the conditions). Another 2 participants were excluded due to Go/NoGo performance and 2 more due to low memory performance.

Pupil data acquisition. The diameter of the left pupil was measured using an EyeLink 1000 System (SR Research), sampled at 1000 Hz. Participants were sat in a darkened room, and asked to maintain fixation whenever possible. A chin rest was used to minimize movement. Iso-luminance was ensured for the grayscale objects and for the blue and yellow frames separately using the Shine toolbox (http://www.mapageweb.umontreal.ca/gosselif/shine)^8^. Stimuli were displayed with 20 degrees of visual angle at a viewing distance of 70 cm. An analogic card of the EyeLink system was connected to an Analogic/Digital converter Cambridge Electronic Device (CED) and data acquired using Spike2 (CED) software. The data were subsequently exported in MATLAB (Mathworks, Natick, MA, USA) format and analyzed using the fieldtrip toolbox (http://www.fieldtriptoolbox.org/).

Pupil data analysis. Pupil diameter data were band pass filtered (0.05 to 4Hz) using a 3rd order Butterworth filter. Blinks were manually detected and corrected by cubic spline interpolation of samples 100 ms either side of the blink. Subsequently, visual artifact rejection was performed to remove bad interpolations. Condition-specific pupil diameter modulations were analyzed using a GLM approach as implemented previously^9^. To dissociate pupil variations due to cognitive processes *vs*. light-evoked pupil constrictions evoked by the appearance of visual stimuli, we modeled these two response components separately. We assumed the pupil to be a linear temporal invariant system (LTI) with an impulse response function for pupillary dilation (*Diameter*(*t*)) described as an Erlang gamma function^10^ which follows this equation as a function of time t:

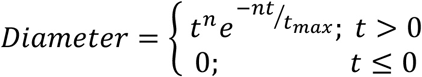

where n and *t*_*max*_ are parameters describing the number of layers of the system and gamma peak time, respectively. To model the cognitive pupillary response we used n = 10.1 and *t*_*max*_ = 0.93s, the parameters estimated in Hoeks and Levelt ^10^. To model the light-evoked visual response, the participant-specific pupil response to encoded stimuli evoking a subsequent familiar (K) judgment at recognition (*i.e*., those trials that were not included in the GLM analysis), were employed as canonical responses to estimate the parameters of interest n and *t*_*max*_ (using the fmincon function in MATLAB) for each participant separately (Figure 3D).

Event-specific responses for our effects of interest (Go and NoGo remembered and forgotten trials) were modeled by convolving a delta-function with the two Erlang gamma basis functions. Nuisance regressors included: the first and second derivatives of these regressors; Go and NoGo familiar trials (convolved with the visual and cognitive response function); fixation cross presentation (convolved with visual response function); button-press reaction time events (convolved with the cognitive response function). The glmfit function in MATLAB was used to calculate parameter estimates for the observed (z-scored) pupil response. For each participant, the ensuing parameter estimates for the cognitive Erlang function were entered into a repeated measures ANOVA to test for a response (Go, NoGo) by subsequent memory (Remembered, Forgotten) interaction across participants.

Lastly, in order to measure the pupil constriction due to the light reflex, band-pass filtered, eye blink-corrected data were epoched into trials, baseline corrected and averaged across all trials for each participant. The maximal pupil constriction (minimum diameter) for each participant was then calculated, and entered into a Welch’s *t*-test (due to different sample sizes) comparing participants who show enhanced memory for Go relative to NoGo stimuli *vs*. those who do not.

### Exp 7

Twenty-three participants (15 females; 23 right handed; age range, 18-35 years; mean age, 30.79; SD, 6.75) performed this experiment. The same presentation time and ISI as in Exp 4-6 were used for both encoding and recognition phases. One subject was excluded from further analysis on the basis of Go/NoGo performance and a further subject due to poor memory performance.

## DATA AVAILABILITY STATEMENT

Methods, including statements of data availability and any associated accession codes and references, are available from the corresponding author on reasonable request

